# Shifted balance between ventral striatal prodynorphin and proenkephalin biases development of cocaine place avoidance

**DOI:** 10.1101/2025.02.28.640871

**Authors:** Amélia Nicot, Pavankumar Yecham, Ilana Serin, David J Barker, Lauren K Dobbs

## Abstract

Evidence from human self-report and rodent models indicate cocaine can induce a negative affective state marked by panic and anxiety, which may reduce future cocaine use or promote co-use with opiates. Dynorphin-mediated signaling within the striatum is associated with negative affect following cocaine withdrawal and stress-induced cocaine seeking. Here, we used a trace conditioning procedure to first establish the optimum parameters to capture this transient cocaine negative affective state in wild type mice, then we investigated striatal opioid peptides as a substrate mediating cocaine conditioned place avoidance (CPA). Previous reports indicate that trace conditioning, where drug administration occurs after removal from the conditioning chamber, results in CPA to ethanol, nicotine, and amphetamine. We tested different cocaine doses, conditioning session lengths, and apparatus types, to determine which combination yields the best cocaine CPA. Cocaine CPA was moderately larger at the highest cocaine dose (25 mg/kg), but this did not generalize across apparatus types and the effect was transient, thus data were collapsed across all parameters. Cocaine conditioning scores were variable, but also became more polarized across conditioning, with approximately equal proportions developing preference and avoidance. We then correlated cocaine CPA with striatal gene expression levels of the opioid peptides enkephalin (*Penk*) and dynorphin (*Pdyn*) using qPCR. Cocaine CPA was correlated with low *Pdyn* levels and a low *Pdyn*:*Penk* ratio in the ventral, but not dorsal, striatum. Consistent with this, mice with higher striatal *Pdyn* relative to *Penk* were more resistant to developing cocaine CPA compared to littermate controls. This approach revealed a subset of subjects sensitive to the aversive state immediately following cocaine administration. Our findings suggest striatal dynorphin has opposing roles in mediating the aversion associated with acute cocaine intoxication versus cocaine withdrawal.

## 1 Introduction

Recent reports indicate the rate of cocaine use in the United States is rising (Orndorff et al., 2024); however, there are no FDA-approved pharmacotherapies and existing off-label medications have limited treatment success (Brandt et al., 2021). Compulsive seeking and taking are hallmarks of cocaine use disorder, and these behaviors are driven by a combination of positive and negative reinforcement mechanisms. While cocaine-induced feelings of euphoria contribute to positive reinforcement, withdrawal from cocaine can provoke an aversive affective state that drives continued cocaine seeking and taking (Gawin and Kleber, 1986; Ettenberg et al., 1999; Sinha et al., 2000; Jhou et al., 2013; Barker et al., 2014). In contrast, acute cocaine intoxication can also induce an aversive affective state, which manifests as a sympathomimetic response, including feelings of paranoia and panic attacks (Anthony et al., 1989; Williamson et al., 1997; Kalayasiri et al., 2006). Epidemiological data indicate cocaine use increases the risk of experiencing panic attacks (Anthony et al., 1989). While this may only occur in a subset of individuals, these negative experiences often lead to decreased cocaine use or stopping altogether (Williamson et al., 1997). Taken together, it appears acute cocaine intoxication can induce bivalent affective states associated with reward and aversion, which generate motivational drives to either seek or avoid cocaine. Accordingly, developing a better understanding of the neurobiology underlying the balance between the rewarding and aversive aspects of acute cocaine intoxication will provide insight into vulnerability and protective factors associated with risk to develop cocaine use disorder and may reveal new targets that can be leveraged for treatment.

While numerous rodent models exist of the positive reinforcing and rewarding aspects of cocaine, including conditioned place preference and operant self-administration, there is limited information on how to best model the aversion associated with acute cocaine use. Acute cocaine has been reported to induce thigmotaxis and decreased time spent in the open arms of an elevated plus maze, suggesting it can induce an aversive state in rodents (Rogerio and Takahashi, 1992; Yang et al., 1992; Simon et al., 1994). In a drug self-administration runway procedure, rats develop an approach-avoidance conflict to a cocaine-associated goal box over progressive training trials (Ettenberg and Geist, 1991, p. 199; Geist and Ettenberg, 1997; Ettenberg, 2004; Jhou et al., 2013; Li et al., 2021; Parrilla-Carrero et al., 2021). In this procedure, subjects traverse a straight-alley runway to receive intravenous cocaine (0.75 – 3.75 mg/kg) when they reach the goal box, and then remain in the goal box for 5 minutes. During this time, the animal associates the goal box with cocaine’s rewarding and aversive properties. Thus, although subjects are in an undrugged state while traversing the runway, over successive training trials, subjects condition an association between the goal box and the drugged state, which results in the expression of approach-avoidance behavior during locomotion to the goal box on subsequent trials. However, with this procedure it is difficult to parse whether the avoidance that develops is the result of cocaine intoxication-associated or cocaine withdrawal-associated aversion.

Place conditioning is a commonly used paradigm to model motivational effects of drugs in experimental animals (see reviews by, Tzschentke, 1998, 2007). The rewarding and aversive effects of drugs (the unconditioned stimulus, US) are inferred based on how much time the subject spends on a drug paired floor (the conditioned stimulus, CS). For instance, simultaneous pairing of the CS and US typically leads to development of preference for a wide range of drugs of abuse, including cocaine, ethanol, morphine, amphetamine, and methamphetamine (Cunningham et al., 1992; Bardo et al., 1995; Dobbs and Cunningham, 2014; Dai et al., 2022). Specifically, when cocaine is administered 0 minutes (i.e., simultaneous pairing) or 5 minutes before being placed in the conditioning chamber, a conditioned place preference (CPP) develops, likely as a consequence of pairing the US with ‘peak or near peak brain levels’ of cocaine (Ettenberg et al., 1999). Conversely, when cocaine is administered 15 minutes before being placed in the conditioning apparatus, a conditioned place avoidance (CPA) develops. Based on the pharmacokinetics of cocaine, this is thought to arise from pairing the aversive aspect of rapidly falling brain levels of cocaine with the conditioning chamber (Ettenberg, 1999). In addition to CPA developing as a consequence of drug withdrawal, several reports indicate the acutely intoxicating effects of drugs of abuse, including ethanol, nicotine, and amphetamine, can be aversive (Fudala and Iwamoto, 1987, 1990; Cunningham and Okorn, 1997). In this conditioning procedure, exposure to the CS precedes the US (Purdy et al., 2001). Rodents are confined to one side of a conditioning apparatus for a specific length of time and then removed, immediately administered a drug injection, and returned to their home cage. This CS-US pairing order is referred to as “trace”, or “delay”, conditioning and is thought to pair a transient, drug-induced aversive affective state with the preceding context. In support of this, trace conditioning with vehicle does not produce a CPA (Fudala and Iwamoto, 1987, 1990), and CPA does not develop when a delay is introduced between the context exposure and ethanol administration (Cunningham and Okorn, 1997). This suggests that the drug imbues the preceding context with motivational significance. Further, the doses of ethanol, nicotine, and amphetamine that produce CPA in a trace conditioning procedure are the same as doses that condition place preference when the CS and US are simultaneously paired, suggesting these different conditioning modalities capture distinct, bivalent affective states of these drugs.

Compared to our understanding of the neurobiological mechanisms underlying cocaine reward, the mechanisms driving cocaine aversion remain relatively uncharacterized. Moreover, much of the knowledge regarding cocaine aversion has focused on mechanisms associated with reinstatement of cocaine seeking following short (4 days) or protracted (up to 30 days) cocaine withdrawal, as opposed to the acutely aversive state induced immediately after cocaine administration. In particular, there is substantial evidence for a role of kappa opioid receptor (KOR) activation by the opioid peptide dynorphin in the aversion associated with repeated cocaine treatment and cocaine withdrawal. Expression levels of prodynorphin (*Pdyn*), the gene that encodes dynorphin, increases in the dorsal striatum following repeated, but not acute, cocaine treatment (Steiner and Gerfen, 1993). This increased dynorphin tone appears to be an adaptive response to repeated activation of D1-receptor containing MSNs (for a review see, Steiner and Gerfen, 1998). Increased dynorphin-mediated KOR signaling also appears to act as a negative reinforcer and increase cocaine seeking. Selective KOR agonists reinstate cocaine seeking after one month of cocaine withdrawal during extinction training in an operant procedure (Valdez et al., 2007). Further, stress-induced release of dynorphin reinstates cocaine seeking after extinction, and this has been reported for wide range of cocaine withdrawal times (4 days – 1 month) (Beardsley et al., 2005; Carey et al., 2007; Redila and Chavkin, 2008). Dynorphin-mediated KOR signaling in distinct regions of the ventral striatum is also acutely aversive by itself (Al-Hasani et al., 2015; Pirino et al., 2020), and activation of KORs in the nucleus accumbens core or ventral tegmental area potentiates nicotine conditioned taste aversion, suggesting KORs may exacerbate the acutely aversive aspect of nicotine (Pham et al., 2022). In contrast, the activity of mu-opioid receptors (MOR) in the ventral striatum is critical for conditioning the rewarding properties of cocaine. Systemic and intra ventral striatum administration of MOR antagonists blocks the acquisition and expression of cocaine place preference (Soderman and Unterwald, 2008; Dai et al., 2022; Matsumura et al., 2023). Additionally, enhancing the level of enkephalin, an opioid peptide with high affinity for MORs and delta-opioid receptors (DOR), within the ventral striatum facilitates acquisition of cocaine place preference (Dai et al., 2022). Taken together, these data suggest that within the ventral striatum, dynorphin-mediated activation of KORs may facilitate withdrawal-associated and perhaps even intoxication-associated cocaine aversion, while enkephalin-mediated activation of MORs supports cocaine’s acutely rewarding state.

Despite evidence that cocaine possesses aversive qualities independent of withdrawal, to date there are no published reports of cocaine CPA for the acutely aversive effects of cocaine. Thus, we predicted that these could be modeled and readily captured using a trace conditioning procedure, which has been used to capture the acutely aversive states of several other abused drugs. Therefore, a major goal of this study was to establish the optimum parameters to condition cocaine CPA using trace conditioning. Interstimulus interval, drug dose, conditioning time, and the type of conditioning apparatus have all been shown to impact the development of a conditioned place preference or avoidance (Cunningham and Okorn, 1997; Tzschentke, 1998; Cunningham et al., 1999; Bevins and Cunningham, 2006). In the current study, we systematically varied cocaine dose, conditioning session length, and conditioning chamber design (i.e., 2-chamber versus 3-chamber) to determine the best set of parameters to induce cocaine CPA. We chose to keep the interstimulus interval short across all subjects, so that cocaine is administered immediately after removal from the conditioning apparatus, because this has been shown to produce the best CPA across drug types (Fudala and Iwamoto, 1987; Cunningham and Okorn, 1997). Additionally, we investigated how the balance between dynorphin and enkephalin within the striatum predicts the development of cocaine CPA using trace conditioning. We used quantitative PCR to measure levels of the prodynorphin (*Pdyn*) and proenkephalin (*Penk*) gene and hypothesized that higher *Pdyn* in the ventral striatum would be correlated with cocaine CPA.

## 2 Materials and Methods

### 2.1 Animals

Male and female adult mice between 8 and 20 weeks of age were used for all experiments. To determine the best method to establish cocaine CPA, we used a variety of wildtype mice, which consisted of in-house bred Cre^-^ littermates or floxed mice from our transgenic mouse lines (all are on a C57BL/6J background). We used 103 mice: 41 *Adora2a-Cre^−/−^* (RRID: MMRRC_034744-UCD; Gerfen et al 2013), 2 MOR^f/f^ (Goldsmith et al., 2013), 19 VGat:Cre (RRID: IMSR_JAX:028862), and 41 VGluT2:Cre (RRID: IMSR_JAX:016963) mice. A subset of these (*Adora2a-Cre^-/-^*= 24), along with 6 additional saline-treated MOR^f/f^ mice to be used as controls, were used in a subsequent qPCR experiment to measure *Pdyn* and *Penk* in the striatum. To test whether a shifted balance between striatal enkephalin and dynorphin affects the development of cocaine CPA, we generated mice with a homozygous deletion of *Penk* from striatal dopamine D2 receptor-containing medium spiny neurons (D2-*Penk*KO). This was achieved by crossing *Adora2a-Cre^+/−^* and *Penk^f/f^*mice. We have used the *Adora2a-Cre* line multiple times to selectively target these medium spiny neurons (Dobbs et al., 2016; Lemos et al., 2016), and this genetic cross produces a near complete reduction in enkephalin from striatal neurons (Schurmann and Zimmer, 2010; Matsumura et al., 2023). Furthermore, these D2-*Penk*KO mice do not show alterations in striatal *Pdyn* expression (Matsumura et al., 2023). For these experiments, 17 D2-*Penk*KO and 17 *Adora2a-Cre^−/−^*;*Penk^f/f^*littermate controls were used. Mice were group-housed in a temperature- and humidity-controlled environment under 12:12 h light/dark cycle (lights on at 06:30 or 7:00) with food and water available *ad libitum*. Behavioral experiments were performed during the light cycle. All animal procedures were performed in accordance with guidelines from the Institutional Animal Care and Use Committees of the University of Texas Austin and Rutgers, the State University of New Jersey.

### 2.2 Drugs

Cocaine HCl was dissolved in sterile saline (0.9%) and administered intraperitoneally at a volume of 10 ml/kg. We tested a range of cocaine doses (15, 20, and 25 mg/kg) to determine the best dose to condition CPA. Previous studies report that rewarding doses of ethanol, nicotine, and amphetamine were able to induce CPA using a Trace Conditioning procedure, thus, we selected cocaine doses known to induce place preference in previous experiments from our lab and others (Cunningham et al., 1999; Dai et al., 2022; Matsumura et al., 2023).

### 2.3 Conditioned Place Avoidance

#### 2.3.1 Apparatus

Mice were divided into two groups and received trace conditioning in either a 2-chamber (Med Associates) or 3-chamber (Stoelting) apparatus (Figure 1A). The 2-chamber apparatus consisted of two distinct contexts that differed by their tactile floor cue (Grid and Hole), which were separated by a guillotine door. The 3-chamber apparatus consisted of two contexts discriminated by tactile floor cues (Smooth and Textured) and visual stimuli (vertical white stripes and plain black walls), which were joined by a smaller connecting chamber.

**Figure 1.**
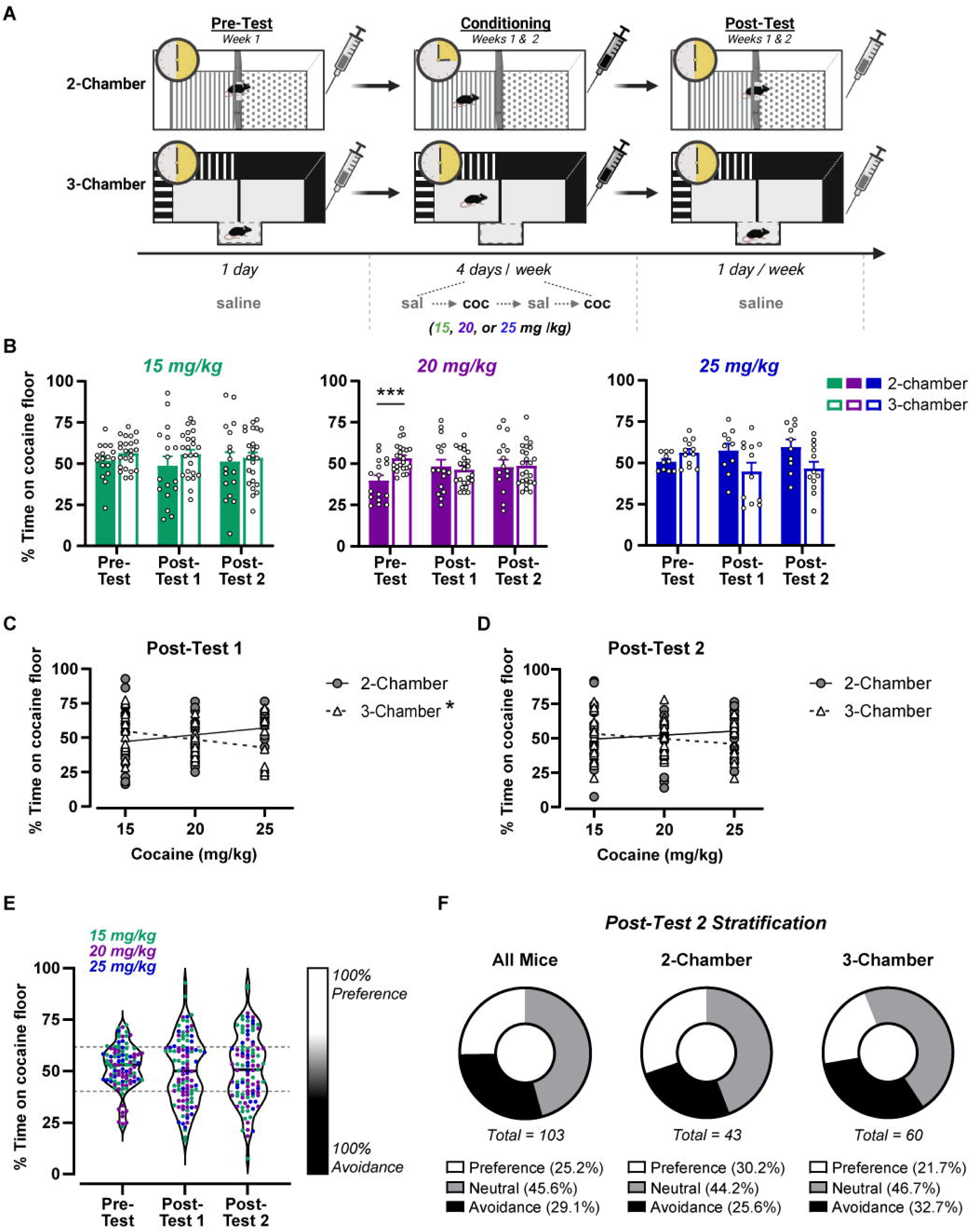
Cocaine dose, apparatus type, and conditioning time do not influence the development of cocaine CPA. (A) Experimental timeline and schematic of conditioning apparatus types for the cocaine trace conditioning procedure. Mice were confined to one side of a 2- or 3-chamber conditioning apparatus for 15- or 30 minutes, respectively. After removal, subjects received cocaine (15, 20, or 25 mg/kg; coc) or saline (sal) injection before returning to their home cage. Cocaine and saline conditioning were alternated over 2 weeks, with preference tests after conditioning days 4 and 8. (**B**) Mean percent time on the cocaine-paired floor during Pretest, Post-Test 1, and Post-Test 2 did not change across tests for any cocaine dose or apparatus type. Pre-Test values were lower for the 2-chamber compared to 3-chamber apparatus before conditioning (20 mg/kg; *** *p* < 0.001). (**C, D**) Linear regressions of cocaine dose versus preference score showed cocaine dose predicts the percent time on the cocaine floor only at Post-Test 1 for the 3-chamber apparatus (* *p* < 0.05). (**E**) Violin plots show distribution of cocaine conditioning when collapsed across cocaine dose and apparatus type. Data were stratified based on one standard deviation of the Pre-Test mean (dashed, grey lines), which created Preference (≥ 61.77%, white), Avoidance (≤ 40.27%, black), and Neutral (40.28 – 61.76%, grey) categories. (F) At Post-Test 2, there were approximately equal proportions of subjects categorized for Preference and Avoidance. These proportions did not significantly differ between apparatus type. Data are expressed as individual data points, mean ± SEM, or a percent of total.

#### 2.3.2 Procedure

##### 2.3.2.1 Experiment 1

To determine the optimum parameters to establish cocaine CPA using Trace Conditioning, we systematically varied cocaine dose, apparatus type, and conditioning session length across two different protocols. Subjects were divided into two conditioning groups and received either 15-minute conditioning sessions in 2-chamber apparatus, or 30-minute conditioning sessions in the 3-chamber apparatus. For both groups, subjects were confined to one side of the conditioning chamber for their specific conditioning time, after which they received an injection of cocaine or saline and were returned to their home cage. Cocaine and saline conditioning sessions were alternated daily. An unbiased subject assignment procedure was used, and the drug-context pairing, drug side, and treatment order were counterbalanced. Subjects received a total of 8 conditioning sessions (4 cocaine and 4 saline) across two weeks. Preference tests were administered prior to conditioning (Pre-Test) and after conditioning sessions 4 and 8 (Post-Test 1 and 2, respectively). Subjects were allowed to freely explore the entire apparatus during preference tests. All preference tests lasted 30 minutes, after which subjects received a saline injection and were returned to their home cage. Place preference and avoidance were determined by calculating the percent time spent on the cocaine paired floor. Different cocaine doses (15, 20, or 25 mg/kg) were tested in separate groups of subjects and for both conditioning groups.

##### 2.3.2.2 Experiment 2

To determine how the balance between striatal dynorphin and enkephalin affect development of cocaine CPA, mice with a selective deletion of enkephalin from striatal D2-MSNs (D2-PenkKO) and littermate controls were trained in a trace conditioning procedure. Subjects were conditioned to 25 mg/kg cocaine using a 2-chamber apparatus with a 15-minute conditioning session length as described in Experiment 1. All procedures and data were calculated as described above.

### 2.4 Quantitative polymerase chain reaction

A subset of mice that had been conditioned with 25 mg/kg cocaine (*Adora2a-Cre^-/--^* = 24) or received equal exposure to repeated saline injection (MOR^f/f^ = 6) were euthanized immediately after Post-Test 2 to measure *Penk* and *Pdyn* using qPCR. Mice were deeply anesthetized with isoflurane, decapitated, the brain extracted and dissected on ice. Tissue was homogenized and stored in RNA Later until RNA was extracted (RNeasy, Qiagen) and cDNA was synthesized (iScript, BioRad). Relative mRNA expression of proenkephalin (*Penk*, Mm01212875_m1) and prodynorphin (*Pdyn*, Mm00457573_m1) was determined relative to the housekeeping gene β-actin (*Actb*, Mm01205647_g1) using TaqMan Gene Expression Assays (ThermoFisher) utilizing a CFX384 Real-Time System (initial hold at 95°C for 20 s, 40 cycles at 95°C for 1 s and 60°C for 20 s). Samples were run in triplicate, and negative controls were run in parallel. Relative expression of the genes of interest was calculated using the ΔΔCт method.

### 2.5 Statistical Analysis

Place conditioning data from the wild-type dataset were quantified as the percent time spent on the cocaine paired floor and analyzed using multifactorial ANOVAs with preference test as a repeated measure and chamber type and cocaine dose as between-subjects factors. To determine whether time spent in the neutral zone of the 3-chamber apparatus was driving place conditioning effects, separate 2-way ANOVAs were performed for each cocaine dose with zone as a between-subjects factor and test as a repeated measure. Sex effects were evaluated using 2-way ANOVA after collapsing place conditioning data across cocaine dose and chamber type. A simple linear regression was used to predict the effect of cocaine dose on cocaine avoidance. Place conditioning datasets for both experiments (wild-type and D2-*Penk*KO) were stratified into preference, avoidance, and neutral based on 1 standard deviation from the respective experiment’s Pre-Test mean. Relative proportions of these groups were analyzed using Chi-square analysis or Fisher’s exact test when sample sizes were below five per group. The relationship between *Penk* and *Pdyn* mRNA expression with place conditioning was analyzed using Pearson’s correlation. Separate 2-way repeated measures ANOVA for preference grouping and opioid peptide gene were performed for the ventral and dorsal striatum. To analyze differences in cocaine avoidance in the D2-*Penk*KO experiment and address the kurtosis observed in the dataset, a Generalized Linear Model with an inverse gaussian probability distribution was used to compare conditioning at Pre-Test versus Post-Test after 2 weeks of conditioning. Significant interactions from multifactorial datasets were followed up with 2-way ANOVA, simple main effects analyses, and pairwise post-hoc t-tests when appropriate. Post-hoc analyses were corrected for multiple comparisons using Sidak’s. Data were analyzed using Prism (GraphPad) or SPSS (IBM), and results were considered significant at an alpha of 0.05. All data are presented are individual values, percentages of total, or mean ± SEM.

## 3 Results

### 3.1 Individual differences in the development of cocaine CPA are not predicted by cocaine dose or apparatus type

We first sought to determine the optimum parameters to establish cocaine CPA using a trace conditioning procedure. We varied the apparatus type (3-chamber or 2-chamber), conditioning time (15-minute or 30-minute), and cocaine dose (15, 20, or 25 mg/kg) between subjects (Figure 1A) and performed a 3-way RMANOVA to determine how these parameters affected the development of cocaine CPA. Overall, the mean percent time spent on the cocaine-paired floor was fairly stable, and we found only modest effects of cocaine dose and apparatus over testing (Test x Chamber x Dose: F_4,230_ = 2.14, *p* = 0.077; Figure 1B). Although there was an interaction between chamber and test, (F_2,230_ = 7.1, *p* < 0.01), follow-up post-hoc analysis shows this was driven by a difference at the Pre-Test, rather than differential conditioning between the chamber types per se (2-chamber vs. 3-chamber at Pre-Test: t_357_ = 3.98, *p* < 0.001). Importantly, this Pre-Test difference appears to be isolated to the 20 mg/kg cocaine dose, suggesting this pre-conditioning bias is a result of sampling error rather than the 2-chamber apparatus being generally biased (Figure 1B, middle). Cocaine dose also affected the percent time spent on the cocaine-paired floor (Main effect of dose: F_2,115_ = 3.43, *p* < 0.05); however, post-hoc analysis did not find significant differences between doses. To further probe this effect, we compared the linear regressions of cocaine dose versus preference score between the two chamber types at Post-Tests 1 and 2. Cocaine dose predicted the percent time spent on the cocaine floor at Post-Test 1 for the 3-chamber apparatus (F_1,58_ = 6.6, *p* < 0.05), but not the 2-chamber (F_1,40_ = 1.76, *p* = 0.19; Figure 1C). A higher cocaine dose was associated with greater avoidance of the cocaine-paired floor in the 3-chamber apparatus; however, this relationship was small, and cocaine dose only accounted for 10% of the variation in preference (*R^2^*= 0.10). In contrast, cocaine dose was not predictive of percent time spent on the cocaine-paired floor at Post-Test 2 for either apparatus type (Figure 1D). Thus, it appears that the highest cocaine dose (25 mg/kg) in the 3-chamber apparatus was slightly better at initially conditioning CPA. Importantly, place avoidance was not due to increased time spent in the non-conditioned “neutral” zone (i.e., the small chamber connecting the two conditioning chambers), but rather less time spent in the cocaine zone and more time spent in the saline zone (Test x Zone: F_4,66_ = 5.7, *p* < 0.001; Table 1). Despite CPA being slightly stronger with this cocaine dose and the 3-chamber apparatus at Post-Test 1, with more conditioning trials neither cocaine dose nor apparatus type influenced expression of CPA.

**Table 1.**
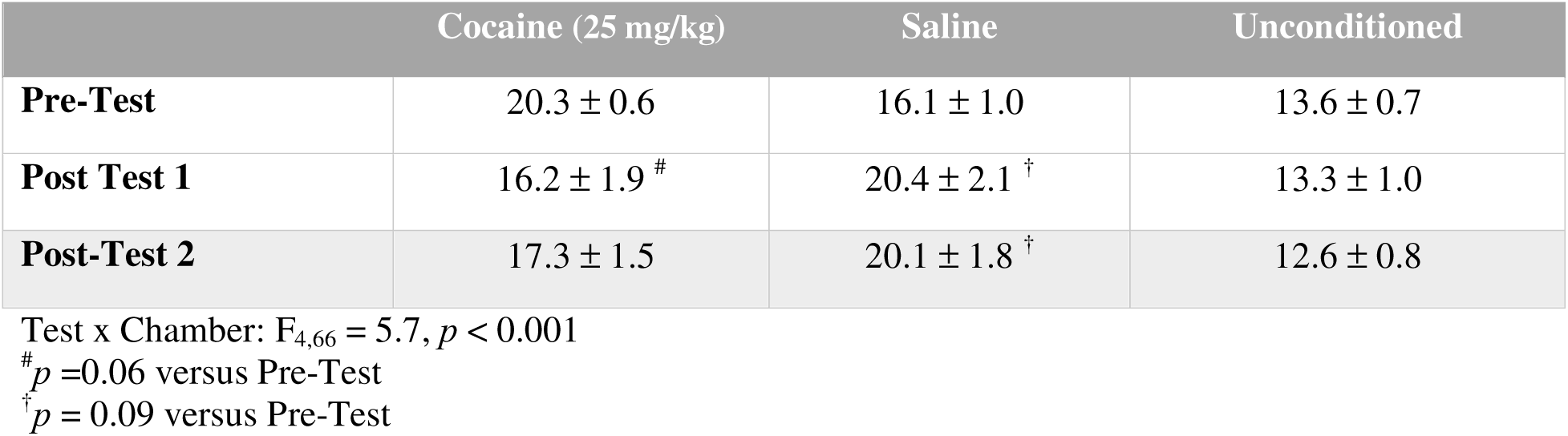
Percent time spent per chamber at each preference test for 25 mg/kg cocaine conditioning.

Given the minimal effect of cocaine dose and apparatus type on the development of cocaine CPA, we collapsed the data across these variables to increase our power to detect sex differences in cocaine CPA (Figure 1E). Cocaine preference did not differ between males and females at any preference test (F_1,95_ = 1.34, *p* = 0.25) and sex did not interact with cocaine dose (F_2,190_ = 1.00, *p* = 0.37), thus all subsequent data analyses were performed collapsed across sex. When analyzing the collapsed data, the mean percent time on the cocaine floor did not change across testing (F_2,204_ = 0.49, *p* = 0.61); however, the variability of the preference scores progressively increased between the Pre-Test and Post-Tests. This was reflected as a greater coefficient of variation at Post-Test 2 (32.5%) compared to Pre-Test (20.8%) and can be observed through the increasingly polarizing responses in Figure 1E. We next examined the relative proportions of subjects that developed a preference or avoidance at Post-Test 2 by stratifying subjects’ Post-Test 2 data based on one standard deviation (σ= 10.75) of the Pre-Test mean. This created 3 categories of subjects: Preference (≥ 61.77 %), Avoidance (≤ 40.27 %), and Neutral (40.28 – 61.76 %; Figure 1F). When evaluating all mice (i.e., collapsed across apparatus type), there was a significant shift in the relative proportions these categories between Pre-Test and Post-Test 2 (Χ^2^ (2, *n* = 103) = 51.08, *p* < 0.0001; Figure 1F and Supplemental Figure 1). Further, the development of cocaine preference or avoidance did not appear to be merely an amplification of the subject’s Pre-Test conditioning score (Supplemental Figure 2). The percentage of subjects expressing avoidance increased by three-fold (from 9.7% to 29.1%). Although the percentage of preferers also increased, it only doubled between Pre-test and Test 2 (from 12.6% to 25.2%). Additionally, the relative proportions of preferers and avoiders was not different between apparatus type, suggesting both apparatus types were equally good at conditioning cocaine CPA (Χ^2^ (2, *n* = 103) = 1.08, *p* = 0.58). When taken together, these data indicate that cocaine causes bivalent responses that become increasingly prevalent at higher doses.

### 3.2 High striatal proenkephalin relative to prodynorphin is associated with cocaine avoidance

We next investigated striatal opioid peptides as an underlying neurobiological mechanism responsible for biasing the development of cocaine preference or avoidance in a trace conditioning procedure. Within the striatum, MOR signaling is linked to motivation for food and cocaine (Hayward et al., 2006; Soderman and Unterwald, 2008; Shin et al., 2010). In particular, augmenting enkephalin within the ventral striatum facilitates acquisition of cocaine place preference (Dai et al., 2022). Conversely, dynorphin-mediated signaling through the kappa opioid receptor is aversive, though there is heterogeneity in this effect depending on the striatal sub-region (Al-Hasani et al., 2015; Pirino et al., 2020). We hypothesized that differential levels of striatal enkephalin relative to dynorphin may be a neurobiological correlate underlying the individual differences in the development of cocaine preference or avoidance following trace conditioning. To test this, we performed RT-qPCR to measure *Penk* and *Pdyn* mRNA levels within the dorsal and ventral striatum of subjects following their completion of trace conditioning (25 mg/kg; 2-chamber apparatus; Figure 2A). We then compared the mRNA expression with each subject’s percent time on the cocaine floor to determine if low *Penk* and high *Pdyn* were associated with cocaine CPA. Cocaine avoidance was not correlated with *Penk* mRNA levels in the dorsal or ventral striatum (Table 2). In contrast, *Pdyn* expression was positively associated with cocaine preference in the ventral, but not dorsal, striatum (*R*^2^ = 0.2, *p* < 0.05; Table 2).

**Figure 2:**
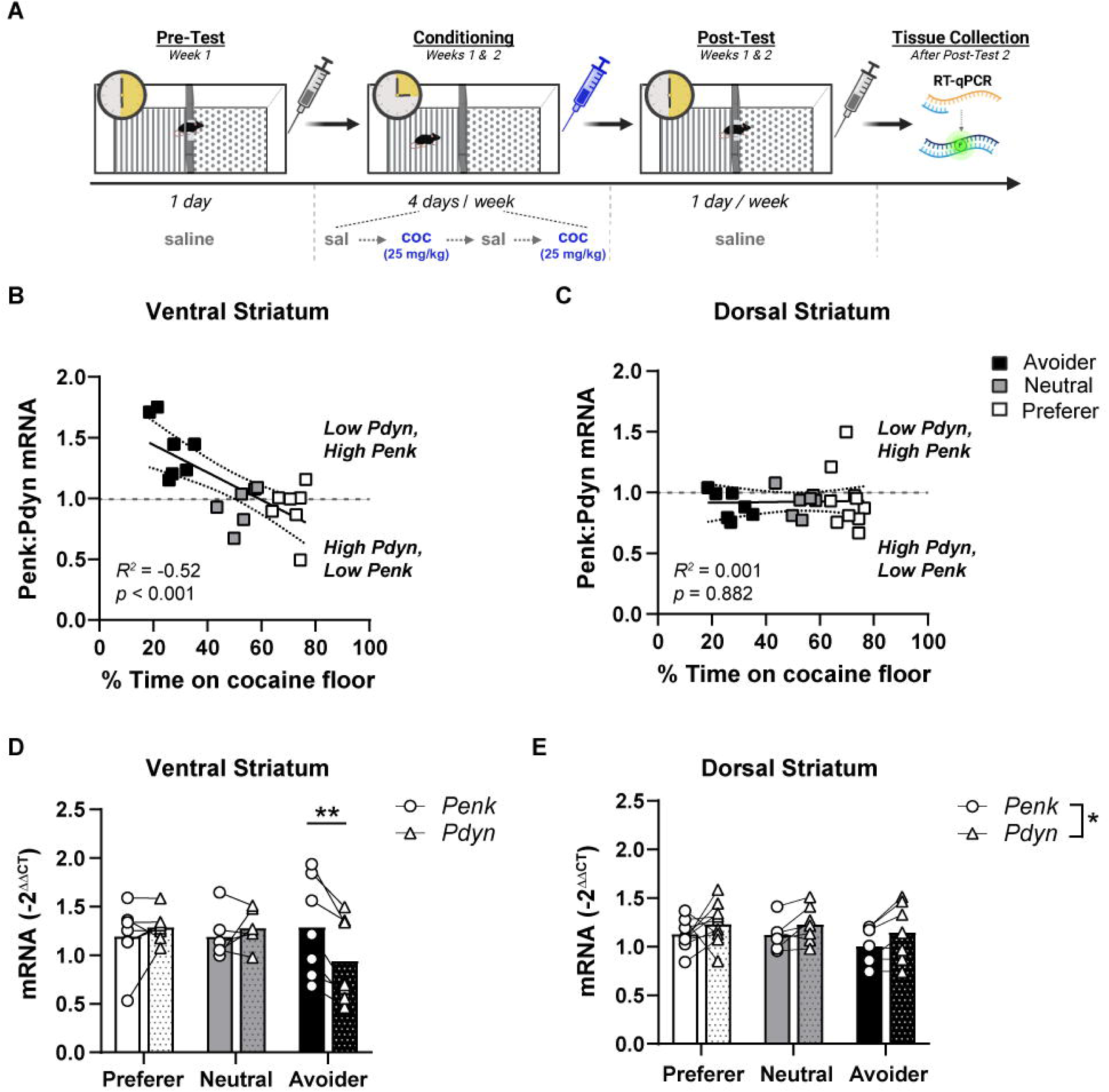
Cocaine CPA is associated with low prodynorphin levels in the ventral striatum. (**A**) Experimental timeline and apparatus schematic for cocaine trace conditioning. Mice were conditioned over two weeks in a 2-chamber apparatus for 15-minutes with 25 mg/kg cocaine. Cocaine or saline was administered after removal from the conditioning chamber. Subjects were euthanized immediately following Post-Test 2 and tissue processed for RT-qPCR. (**B, C**) The ratio of *Penk*:*Pdyn* mRNA was negatively correlated with the percent time spent on the cocaine paired floor in the ventral (**B**), but not dorsal (**C**), striatum. Subjects expressing cocaine avoidance (black), neutral conditioning (grey), or preference (white) were stratified based on one standard deviation from the Pre-Test mean (see Figure 1). (**D**) Subjects categorized as Avoiders (black bars), but not Preferers (white bars) or Neutral subjects (grey bars) had lower *Pdyn* mRNA compared to *Penk* levels in the ventral striatum (Gene x Group interaction: F_2, 17_ = 9.7, *p* < 0.01; post-hoc t-test: ** *p* < 0.01). (**E**) *Pdyn* was slightly higher than *Penk* levels for Preferers, Avoiders, and Neutral subjects in the dorsal striatum (Gene main effect, * *p* < 0.05). Data are expressed as individual data points or the mean.

**Table 2.** Correlations (*R^2^*) of percent time on the cocaine floor with striatal *Penk* and *Pdyn* mRNA.

|  | *Penk*, DS | *Penk*, VS | *Pdyn*, DS | *Pdyn*, VS |
| --- | --- | --- | --- | --- |
| **% time on cocaine floor** | 0.095 | 0.0001 | 0.055 | 0.2004* |

\* $p < 0.05$
DS, dorsal striatum
VS, ventral striatum

A correlation between cocaine preference and *Pydn*, but not *Penk*, suggests a shift in the balance between these peptides may be influencing development of cocaine preference versus avoidance. Indeed, dynorphin and enkephalin regulate the responsiveness of striatal medium spiny neurons and in this way play an important role in regulating striatal output, motor activity, and motivated behavior (Steiner & Gerfen, 1998). Thus, we correlated the *Penk*-to-*Pdyn* ratio with each subject’s cocaine preference. Within the ventral striatum, a higher *Penk*:*Pdyn* was negatively correlated with percent time spent on the cocaine floor (*R^2^* = -0.52, *p* < 0.001; Figure 2B). In contrast, cocaine preference was not correlated with the *Penk*:*Pdyn* ratio in the dorsal striatum (Figure 2D).

The strong negative association between ventral striatum *Penk*:*Pdyn* with cocaine preference indicates the balance of these peptides may be important for regulating cocaine avoidance. However, this analysis cannot ascertain whether this is due to higher *Penk*, lower *Pdyn*, or a combination of the two. To address this, we stratified each subject’s mRNA expression into Preference, Avoidance and Neutral groups using 1 standard-deviation of the Pre-Test mean (Figure 1E). Within the ventral striatum, the relationship between *Penk* and *Pdyn* mRNA expression depended on preference group (Gene x Group: F_2,17_ = 9.7, *p* < 0.01; Figure 2D). Subjects categorized as avoiders had higher *Penk* levels relative to *Pdyn* (t_17_ = 4.32, *p* < 0.01). In contrast, there was no difference between *Penk* and *Pdyn* for subjects that preferred cocaine or were neutral. Additionally, while mean levels of *Pdyn* were lower for avoiders compared to preferers or neutral subjects, this comparison did not reach statistical significance. Within the dorsal striatum, the balance between *Penk* and *Pdyn* was similar across avoiders, preferers, and neutral subjects, with *Pdyn* being slightly elevated relative to *Penk* across all groups (Gene: F_1,21_ = 7.03, *p* < 0.05; Figure 2E). Taken together, cocaine CPA is associated with lower *Pdyn* levels relative to *Penk* selectively in the ventral striatum.

### 3.3 Higher relative Pdyn within the striatum prevents development of cocaine avoidance

While the qPCR results are informative, they cannot preclude whether differences in *Pdyn* and *Penk* existed prior to conditioning. Thus, we next tested whether the lower *Pdyn* relative to *Penk* levels in the ventral striatum of avoiders represented a causal, pre-existing neural signature that biases towards development of CPA in a trace conditioning procedure. To address this, we took advantage of a cell-selective knockout mouse line in our laboratory that lacks *Penk* in striatal D2-MSNs (D2-*Penk*KO; Matsumura et al., 2023). These mice show a near complete lack of *Penk* in the striatum (Matsumura et al., 2023) and therefore have a pre-existing shift in the striatal *Penk*:*Pdyn* ratio that favors a higher *Pdyn* tone relative to *Penk*. Given our observations that, 1) higher *Penk* relative to *Pdyn* is associated with cocaine avoidance, and 2) higher *Pdyn* relative to *Penk* is associated with cocaine preference, we hypothesized that D2-*Penk*KO mice would be more resistant to developing cocaine CPA. Additionally, since wild-type mice show a wide distribution of preferers versus avoiders, we speculated that a difference between the D2-*Penk*KOs and littermate controls would likely be best reflected as a shift in the relative proportions of preferers, avoiders, and neutral mice rather than in differences in the mean percent time on the cocaine floor per se.

The distribution of preference scores in the D2-*Penk*KO and littermate controls was not normally distributed and exhibited significant kurtosis for Pre-Test and Post-Test 2. Therefore, we used a Generalized Linear Model (Wald’s Chi square) with an inverse Gaussian distribution to approximate the distribution and analyze the change in preference between Pre-Test and Post-Test 2. We observed that the percent time on the cocaine floor changed across testing depending on the genotype (Genotype x Test: Wald Chi-Square: 5.73, df = 1, *p* < 0.05; Figure 3B-C), despite no difference between genotype in the pre-test (Figure 3C). Follow-up analysis confirmed that by Post-Test 2, *Penk^f/f^* controls had developed a significant cocaine avoidance (*p* < 0.05) compared to the Pre-Test. In contrast, cocaine CPA was mitigated in D2-PenkKOs (*p* = 0.12) (Figure 3C). Moreover, the difference between genotypes was significant (*p* < 0.05), suggesting that a higher *Pdyn* tone relative to *Penk* prevented the development of cocaine avoidance. This finding was confirmed by further evaluation of the data after stratifying subjects into Preference, Avoidance, and Neutral groups based on one standard deviation from the mean preference at Pre-Test (*X̄*= 45.14, σ = 11.84). The mean conditioning score for *Penk^f/f^* controls (36.5%), but not for D2-*Penk*KOs (52.4%), fell inside the region of avoidance (< 38.5%). At Pre-Test, the relative proportions of Preference, Avoidance, and Neutral were not different between genotypes, and most subjects fell into the neutral range (Fisher’s exact test; *p* = 0.69; Figure 3D). I At Post-Test 2, both genotypes showed a significant shift in the relative proportions of these groups (Fisher’s exact test; *Penk^f/f^*: *p* < 0.0001; D2-*Penk*KO: *p* < 0.01); however, they were not different from one another (Fisher’s exact test; *p* = 0.12; Figure 3F). Since the inherently large variation in this kind of dataset coupled with a small sample size when stratified into three groups may have underpowered this analysis, we re-stratified groups into a binary classification, Avoidance and Non-avoidance. At Pre-Test, most subjects were categorized as Non-avoidance and there was no difference between genotypes (Χ*^2^* (1, *N* = 34) = 0.4, *p* = 0.5; Figure 3E). After conditioning, at Post-Test 2, nearly half as many D2-*Penk*KOs developed avoidance compared to *Penk^f/f^* controls (35.3% vs. 64.7%), though this difference only approached significance (Χ*^2^* (1, *N* = 34) = 2.9, *p* = 0.08; Figure 3G). When comparing the percentage of Preferers between genotypes, there was no difference at Pre-Test (D2-*Penk*KOs: 8.82%, *Penk*^f/f^: 2.94%; Fishers exact test: *p* = 0.60). At Post-Test 2, the percentage of Preferers increased for both genotypes, but neither increase was significant (D2-*Penk*KO: 23.5%, *p* = 0.14; *Penk*^f/f^: 17.65%, *p* = 0.09). Additionally, there was no difference between genotypes in the percentage of Preferers at Post-Test 2 (*p* = 0.73). Taken together, these data suggest that the balance between striatal *Penk* and *Pdyn* may influence development of cocaine preference or avoidance using a trace conditioning procedure.

**Figure 3:**
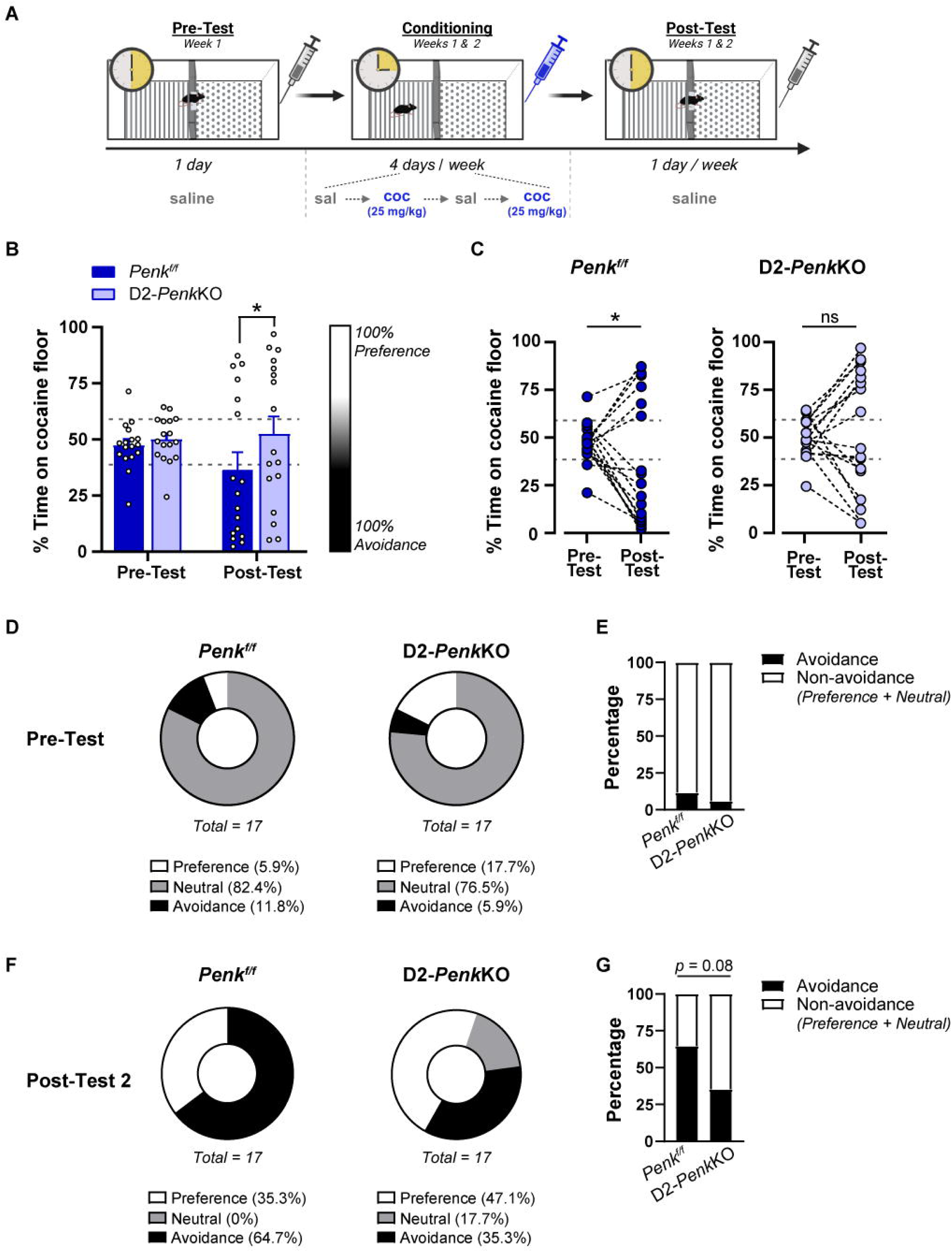
High relative striatal prodynorphin prevents development of cocaine CPA. **(A**) Experimental timeline and apparatus schematic for cocaine trace conditioning. Mice lacking proenkephalin from D2-receptor containing medium spiny neurons (D2-*Penk*KO) and littermate controls (*Penk*^f/f^) were conditioned over two weeks in a 2-chamber apparatus for 15-minutes with 25 mg/kg cocaine. Cocaine or saline was administered after removal from the conditioning chamber. (**B**) *Penk*^f/f^ subjects had lower percent time spent on the cocaine paired floor compared to D2-*Penk*KOs at the Post-Test (* *p* < 0.05). (**C**) *Penk*^f/f^ developed a cocaine CPA (* *p* < 0.001), while D2-*Penk*KO developed a cocaine place preference (** *p* < 0.01) at Post-Test 2 compared to Pre-Test. (**D, F**) Subjects were stratified into Preference, Neutral, and Avoidance categories based on one standard deviation from Pre-Test Mean (dashed, grey line in panel **B**). Most subjects were categorized at Neutral at Pre-Test (**D**). At Post-Test 2, more *Penk*^f/f^ subjects were categorized as Avoidance than D2-*Penk*KO, but this was not significant. (**E, G**) When stratified into a binary classification of Avoidance (black) and Non-avoidance (white; Preference + Neutral), twice as many *Penk*^f/f^ categorized as Avoidance than D2-*Penk*KOs (# *p* = 0.08). Data are expressed as individual data points, mean ± SEM, or a percent of total.

## 4 Discussion

### 4.1 Optimum parameters for cocaine CPA

In this investigation, we sought to establish the optimum parameters to condition cocaine place avoidance using a trace conditioning procedure and to investigate the striatal opioid milieu as a neurobiological correlate of cocaine CPA. Previous research has shown CPA develops following trace conditioning to ethanol, amphetamines, and nicotine. However, despite self-report and preclinical data indicating cocaine has aversive qualities associated with intoxication, CPA to cocaine using trace conditioning has not been established. Moreover, CPA for cocaine has traditionally proven more difficult in mouse models, as compared to work in rats (Jhou et al., 2013). We tested different cocaine doses, conditioning session times, and apparatus types, which are variables previously shown to influence development of drug conditioning. We observed a large variability in cocaine conditioning, with roughly equal proportions of subjects developing preference and avoidance. Quite to our surprise, the propensity to condition avoidance was not reliably predicted by any of the parameters we manipulated, nor was backwards conditioning more effective (unpublished observations). Additionally, sex was not a factor in determining the likelihood of conditioning cocaine place avoidance. Although conditioning time varied between the 2-chamber and 3-chamber apparatuses, there was no difference between these experimental results, which speaks to the robust nature and generalizability of these findings.

The small effect of cocaine dose on cocaine CPA was specific to the 3-chamber apparatus and did not persist past 1 week of conditioning. It is possible a higher dose (i.e., > 25 mg/kg) may be more acutely aversive and therefore condition a larger, more enduring CPA. Using a simultaneous conditioning procedure, most doses of methamphetamine resulted in CPA, while only very low doses produced conditioned preferences (Cunningham and Noble, 1992). To our knowledge, though, there are no reports showing an inverted-U dose-response for cocaine preference with simultaneous conditioning. Thus, it is unclear whether the balance between reward and aversion shifts toward more aversion with increasing cocaine dose, or perhaps whether the aversive qualities of cocaine require a longer history of drug use. It should be noted, however, that acutely aversive drug doses do not appear to be necessary because previous experiments use rewarding drug doses to induce CPA in trace conditioning (Fudala and Iwamoto, 1987, 1990; Cunningham and Okorn, 1997).

Mean CPA was slightly larger when subjects were conditioned using a 30-minute session time within the 3-chamber apparatus. In simultaneous conditioning procedures, conditioning session time does not affect cocaine place preference (Cunningham et al., 1999). It is conceivable that longer conditioning times may lead to a stronger memory of the CS when the drug is administered, and thus a better CPA; however, the small effect in the current dataset suggests this is not the case. Consistent with this, ethanol CPA develops with only a 5-minute conditioning length (Cunningham and Okorn, 1997). One advantage of the 3-chamber apparatus is the distinct presentation of conditioned cues. However, data from 3-chamber conditioning apparatuses, while commonly used, can also be difficult to interpret if the time spent in the unconditioned “neutral” chamber changes significantly after conditioning. Subjects may spend more time in the unconditioned chamber due to its relative novelty, and drugs that block the habituation to novelty may enhance this effect (Parker, 1992). In the current experiment, time spent in unconditioned chamber did not significantly change after conditioning, indicating that the preference and avoidance observed were not the result of shifts in novelty perception of the unconditioned chamber. Taken together, the trace conditioning procedure yields fairly equal cocaine conditioning between the 2-chamber and 3-chamber apparatuses, indicating neither is preferential for generating cocaine CPA.

### 4.2 Interpretations of cocaine CPA in trace conditioning

Several potential interpretations have been posited to explain the development of drug CPA using trace conditioning, including interfering with habituation of neophobia (Fudala and Iwamoto, 1987), the drug-paired context becoming a conditioned inhibitor potentially by conditioning a “drug wanting” state (Carelli and West, 2014), or by conditioning an initial, transient aversive drug effect (Cunningham and Okorn, 1997). If the drug interferes with habituation of neophobia associated with the conditioning chamber, CPA may represent contextual neophobia rather than an aversive aspect of the drug per se. However, this interpretation does not explain why subjects express place preference when the same drug is administered in a simultaneous conditioning procedure. Thus, while we cannot rule this out for other drugs with dissociative properties, this appears to be an unsatisfactory explanation for cocaine CPA in the current study.

In the case of the drug-paired context becoming a Pavlovian conditioned inhibitor, the conditioning chamber is thought to elicit avoidance because it predicts the absence of the rewarding drug sensation and a “drug wanting” state (Wheeler et al., 2008, 2015). However, morphine and diazepam fail to condition a place avoidance (or preference) in a trace conditioning procedure (Kumar, 1972; Sherman et al., 1980; Bardo et al., 1984; Spyraki et al., 1985). This does not appear due to subthreshold dosing because the same doses elicited place preference using a simultaneous conditioning procedure (Bardo et al., 1984; Spyraki et al., 1985). Additionally, if the delay of drug reward was indeed driving the CPA, one would expect the CPA to maintain, and perhaps intensify, with increasing delay between the CS and US. When increasing delays were introduced in ethanol trace conditioning experiments, though, CPA is abolished (Cunningham and Okorn, 1997). In contrast, studies using a conditioned taste aversion procedure report robust aversion with a 30-minute delay between the CS and cocaine self-administration (Wheeler et al., 2008). However, despite similar CS-US temporal dynamics between trace conditioning and conditioned taste aversion experiments, the conditioned responses to contextual versus gustatory stimuli can be different. For instance, morphine administration following consumption of a novel tastant within a conditioning chamber, results in taste, but not place, avoidance (Bardo et al., 1984). Notably, stronger relationships between the qualities of specific cues and the subsequent reinforcers have long been shown to support stronger conditioning (Garcia and Koelling, 1966). Thus, it is possible that the ability of a stimulus to support Pavlovian associations with motivational drug effects depends on the stimulus modalities and specific physiologic drug response. However, coupling these conditioning discrepancies with evidence that CPA degrades with increasing delays, makes the conditioned inhibitor explanation of CPA less likely. Moreover, conditioned inhibition does not reasonably explain why in the current study approximately equal proportions of subjects developed preference and avoidance.

The development of CPA in trace conditioning may also result from subjects experiencing a transient aversive state following cocaine injection, which becomes paired with the preceding context. The onset of a longer-lasting rewarding affective state is thought to occur afterwards and is more likely paired with the home cage environment. This transient aversion may stem from various sources such as the transition from sobriety to intoxication, peritoneal irritation, or an aversive state associated with cocaine intoxication. We suspect aversion due to state transition or peritoneal irritation is unlikely. Only a subset of subjects developed CPA in the current study and not all intoxicating drugs condition avoidance. Further, there is not convincing evidence for peritoneal irritation at the cocaine doses used in the current study and, if present, this would occur equally for the saline paired chamber. Evidence from human cocaine users (Anthony et al., 1989; Williamson et al., 1997; Kalayasiri et al., 2006) and rodent models (Rogerio and Takahashi, 1992; Yang et al., 1992; Simon et al., 1994) indicate that cocaine elicits an anxiogenic state immediately after administration in a subset of subjects, supporting the hypothesis that CPA in the current study resulted from an acutely aversive state associated with cocaine intoxication. How long this aversive state lasts is not known, but it is likely a combination of pharmacokinetic and pharmacodynamic effects. Despite this, the culmination of evidence suggests cocaine CPA likely results from a Pavlovian association between a transient, cocaine-induced negative affective state and the preceding context.

### 4.3 Striatal opioids as a mechanism influencing the development of cocaine avoidance

Dynorphin-mediated KOR signaling within the ventral striatum is associated with cocaine withdrawal (Chartoff et al., 2012) and pain-induced negative affect (Massaly et al., 2019), and withdrawal from chronic cocaine increases dynorphin levels within the striatum (Steiner and Gerfen, 1998). This heightened dynorphin tone is thought to act as a negative reinforcer, because KOR activation incites stress-induced reinstatement of cocaine seeking (Carey et al., 2007; Schindler et al., 2012) and blockade of KORs attenuates it (Beardsley et al., 2005; Aldrich et al., 2009). Based on this literature, we hypothesized cocaine CPA would be associated with high striatal dynorphin. Conversely, subjects expressing cocaine CPA had lower levels of striatal *Pdyn* compared with “Preferer” and “Neutral” subjects, and this effect was isolated to the ventral striatum. Together, these data suggest striatal dynorphin signaling may have opposing roles in mediating aversion associated with acute intoxication versus cocaine withdrawal. Considering KOR-mediated signaling facilitates cocaine relapse after withdrawal (Valenza et al., 2020), the association between low striatal *Pdyn* levels and cocaine CPA in the current data support the interpretation that avoidance is due to a transient cocaine aversion, rather than a “drug wanting” state.

The finding that mice with higher relative striatal dynorphin to enkephalin (i.e., D2-*Penk*KO) have greater cocaine seeking is consistent with the literature and supports the hypothesis that high striatal dynorphin tone drives cocaine seeking. Thus, place preference may reflect the experience of an aversive “drug wanting” state mediated by high striatal dynorphin. This is supported by the current data in D2-*Penk*KO mice since they have higher *Pdyn*:*Penk* levels compared to littermate controls. Consistent with this, the non-selective opioid antagonist naloxone blocks expression of cocaine preference in female D2-*Penk*KO mice, but not littermate controls (Matsumura et al., 2023). Whether this effect was mediated by dynorphin or another opioid peptide, though, is unclear. This is an important consideration because enkephalin tone and MOR-mediated signaling within the ventral striatum also facilitate the acquisition and expression of cocaine place preference (Dai et al., 2022). Another potential interpretation is that D2-*Penk*KO mice have a generally potentiated cocaine reward, though our previous work indicates D2-*Penk*KO mice acquire cocaine place preference normally in a simultaneous conditioning procedure (Matsumura et al., 2023). Dynorphin-mediated “drug wanting” may be a less probable explanation for cocaine preference in the wild type mice from the current study because *Pdyn* levels for “Preferers” were similar to that of “Neutral” and “Avoiders”. Thus, KOR-mediated signaling was likely not driving cocaine seeking in these subjects. It is possible place preference is a result of unmasking cocaine reward due to a blunted transient aversion. This is also unlikely, though, since rewarding doses of opiates and benzodiazepines do not condition avoidance or preference in a trace conditioning procedure (Sherman et al., 1980; Bardo et al., 1984; Spyraki et al., 1985), but do result in preference in simultaneous conditioning (Bardo et al., 1984). Taken together these data suggest that while enkephalin and dynorphin both drive cocaine seeking, they do so through opposing motivational mechanisms. Moreover, an imbalance between these endogenous opioids, in which dynorphin levels are low relative to enkephalin, may render subjects more sensitive to the transient aversion associated with cocaine intoxication.

### 4.4 Limitations to the current study

The selectivity of the association between dynorphin and cocaine CPA for the ventral striatum is consistent with previous reports indicating dynorphin signaling specifically within the ventral striatum is associated with cocaine withdrawal. However, work by Al-Hasani and colleagues (Al-Hasani et al., 2015) indicate even further sub-region specificity related to dynorphin-mediated CPA, with KOR signaling within the ventral, but not dorsal, nucleus accumbens shell conditioning place avoidance. Although it is unclear if this subregion distinction is conserved regarding dynorphin’s role in cocaine conditioning, the brain dissections and qPCR used in the current study cannot distinguish between these subregions and thus is a limitation of this approach. Additionally, the use of *Penk* and *Pdyn* mRNA expression analysis to probe differences in the endogenous opioid milieu, while highly quantitative, may not completely correlate with enkephalin and dynorphin peptide levels. The use of semi-quantitative approaches to measure peptide levels, such as immunofluorescence, may complement this approach in future studies. Lastly, it remains unclear whether the qPCR results represent pre-existing differences that lead to CPA, or whether the differences in *Penk* and *Pdyn* mRNA expression developed alongside cocaine conditioning. While we unfortunately cannot sample the brain prior to conditioning, the culmination of our results suggests both interpretations may be possible. The gradual development of CPA and preference may indicate a shift in dynorphin tone occurs because of cocaine exposure. This is consistent with time course studies showing striatal *Pdyn* levels increase in order to restrain cocaine-induced activation of D1-MSNs (Steiner and Gerfen, 1998). However, our finding that D2-PenkKOs are resistant to developing cocaine CPA may also indicate that a pre-existing shift in the balance between striatal dynorphin and enkephalin can predispose development of cocaine CPA. Future experiments capable of directly assessing striatal dynorphin tone in real time or using calcium imaging to track activity of D1- and D2-MSNs will undoubtedly be beneficial in addressing questions as these.

Our findings suggest enkephalin and dynorphin within the striatum are important for expression of conditioned rewarding and aversive aspects of cocaine. It is tempting to speculate that levels of dynorphin itself might be most essential to the expression of place preference or avoidance because “Avoiders” showed lower *Pdyn* levels compared to “Neutral” and “Preferer” mice, with no differences in *Penk* levels. However, because shifts in *Pdyn* fundamentally alter the balance with *Penk*, these data cannot distinguish whether absolute peptide level or balance between the peptides is responsible for the effect. Similarly, genetically deleting *Penk* from striatal MSNs also inherently shifts the balance to favor dynorphin signaling within the striatum. Future experiments using intracranial administration of short-acting KOR antagonists during preference testing will help disentangle how acute changes in dynorphin-mediated signaling affect expression of CPA without inducing long-term alterations in the enkephalin-dynorphin balance.

When testing different parameters to establish cocaine CPA in Experiment 1, we observed a slightly larger CPA with the 30-minute conditioning session. Though, since this conditioning time was only tested in the 3-chamber apparatus, we cannot differentiate whether the timing or apparatus type was the main driver for this effect. For Experiment 2, we used the highest cocaine dose to establish cocaine CPA. Interestingly, a larger proportion of control mice developed CPA in Experiment 2 than in the initial experiment, which may in part have driven the genotype difference. Considering the smaller sample size in Experiment 2, we cannot rule out the possibility that sampling error contributed to the more robust CPA in controls. While it is tempting to compare the D2-*Penk*KO mice to control mice from Experiment 1, the most appropriate comparison is the littermate controls run in parallel. Additionally, it is not clear whether this genotype difference would persist across other cocaine doses.

### 4.5 Conclusions

Cocaine trace conditioning is a good approach to observe the individual differences in the development of cocaine preference and avoidance and a means to identify subjects that are more vulnerable to the brief aversive state that occurs following the administration of a rewarding cocaine dose. This transient aversive state can curb future cocaine use, suggesting an increased sensitivity to the aversive component may be protective against developing cocaine abuse. Conversely, these individuals may be more prone to co-abuse opioids to counteract this aversive state. The balance between striatal enkephalin and dynorphin may represent a neurobiological mechanism underlying the vulnerability to the transient aversion induced by cocaine. Future experiments can leverage this procedure and these kinds of data to further probe the causal role of striatal opioids in this behavior and identify new mechanisms that are potentially protective against cocaine abuse or even predictive of cocaine-opiate co-abuse.

## Author Contributions

Conceptualization and Methodology: DJB and LKD. Investigation and Formal analysis: DJB, LKD, AN, and PK. Writing – original draft: LKD and AN. Writing – review and editing: DJB, LKD, and AN. Supervision, Funding acquisition, and Resources: DJB and LKD.

## Ethics Statement

All animal procedures were performed in accordance with the guidelines from the Institutional Animal Care and Use Committees of the University of Texas Austin and Rutgers, the State University of New Jersey.

## Data Availability Statement

The data supporting the findings of this study are available on reasonable request from the corresponding author.

## Acknowledgments

We would like to thank Harshvir Kaur for excellent assistance in running the three-chamber CPA experiments, and Jennifer Mejaes for her guidance during these experiments.

