## Supplemental Figure 1 for "Shifted balance between ventral striatal prodynorphin and proenkephalin biases development of cocaine place avoidance"

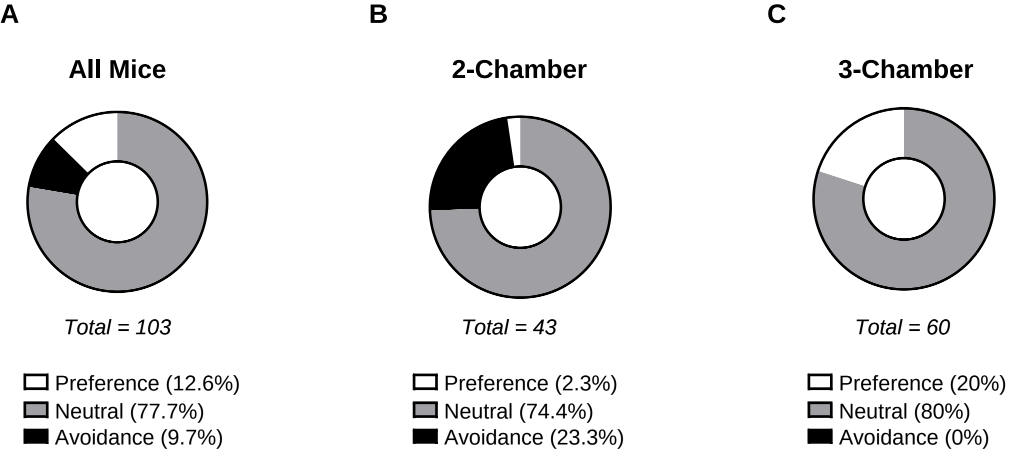


**Supplemental Figure 1. Relative proportions of pre-conditioning Preference, Neutral and Avoidance.** (A-C) Data were stratified based on one standard deviation of the Pre-Test mean (dashed, grey line in Figure 1), which created Preference (≥ 61.77%, white), Avoidance (≤ 40.27%, black), and Neutral (40.28 – 61.76%, grey) categories. Relative proportions of these categories are shown for the total cohort (A), for mice conditioned in the 2-chamber apparatus (B), and in the 3-chamber apparatus (C). Data are expressed as a percent of total. Related to Figure 1.
