## Supplemental Figure 2 for "Shifted balance between ventral striatal prodynorphin and proenkephalin biases development of cocaine place avoidance"

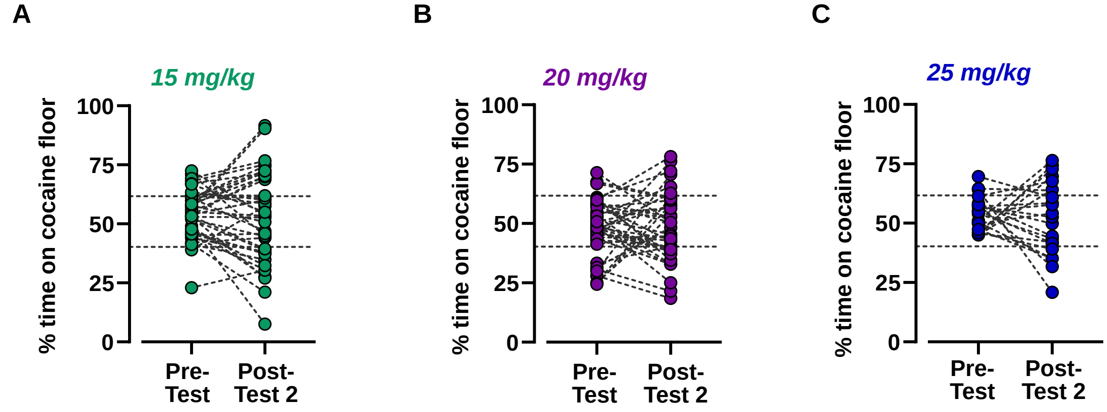


**Supplemental Figure 2. Development of cocaine preference or avoidance is not determined by a pre-conditioning bias.** (A-C) Data were stratified based on one standard deviation of the Pre-Test mean (dashed, grey lines), which created Preference (≥ 61.77%, white), Avoidance (≤ 40.27%, black), and Neutral (40.28 – 61.76%, grey) categories. Conditioning scores at Pre-Test and Post-Test 2 are shown paired for each subject conditioned with 15 mg/kg (A), 20 mg/kg (B), or 25 mg/kg (C) cocaine. Data are expressed as individual values. Related to Figure 1.
